# Rapid expansion of immune-related gene families in the house fly, *Musca domestica*

**DOI:** 10.1101/068213

**Authors:** Timothy B. Sackton, Brian P. Lazzaro, Andrew G. Clark

## Abstract

The house fly, *Musca domestica*, occupies an unusual diversity of potentially septic niches among sequenced Dipteran insects and is a vector of numerous diseases of humans and livestock. In the present study, we apply whole-transcriptome sequencing to identify genes whose expression is regulated in adult flies by bacterial infection. We then combine the transcriptomic data with analysis of rates of gene duplication and loss to provide insight into the evolutionary dynamics of immune-related genes. Genes up-regulated after bacterial infection are biased toward being evolutionarily recent innovations, suggesting the recruitment of novel immune components in the *M. domestica* or ancestral Dipteran lineages. In addition, using new models of gene family evolution, we show that several different classes of immune-related genes, particularly those involved in either pathogen recognition or pathogen killing, are duplicating at a significantly accelerated rate on the *M. domestica* lineage relative to other Dipterans. Taken together, these results suggest that the *M. domestica* immune response includes an unusual diversity of genes, perhaps as a consequence of its lifestyle in septic environments.

## Introduction

The rapid increase in the number of sequenced genomes over the past decade has dramatically reshaped our understanding of the evolutionary dynamics of the insect innate immune system. It has long been recognized that genes involved in the immune response are among the most rapidly evolving in many organisms, including mammals (Hughes and Nei 1988; Nielsen et al. 2005; Kosiol et al. 2008), plants (Tiffin and Moeller 2006), and insects (Sackton et al. 2007; Lazzaro 2008; Obbard et al. 2009), with adaptation presumably driven by host-pathogen conflict. In the era of comparative genomics, it has become clear that this pattern of rapid evolution occurs against a backdrop of deeply conserved orthology in core signaling transduction pathways (Toll, imd, JAK/STAT, and JNK) across most insects studied to date (Evans et al. 2006; Sackton et al. 2007; Waterhouse et al. 2007; Werren et al. 2010),with only rare examples of secondary loss (Gerardo et al. 2010).

In addition to signaling cascades that are activated in response to infection, insect immune systems contain classes of proteins involved in pathogen recognition as well as classes of effector proteins such as antimicrobial peptides whose role is to clear infections. Both recognition proteins and effector proteins are encoded by a diverse array of gene families with a variety of functions and specificities. In contrast, immune signaling tends to occur through only four primary signal transduction pathways – Toll, imd, JAK/STAT, and JNK (Buchon et al. 2014). While components of these primary signal transduction pathways are typically conserved in 1:1 orthology across all insects, gene families encoding recognition or effector proteins often vary considerably in copy number between species and exhibit substantial rates of duplication and deletion within evolutionary lineages (Ghosh et al. 2011). Several gene families, especially those encoding antimicrobial peptides, are restricted to particular insect clades (Bulet et al. 1999; Vizioli et al. 2001; Sackton et al. 2007), and the transcriptional response to infection in at least some insects results in the upregulation of numerous taxonomically-restricted genes (Sackton et al. 2013).

The house fly, *Musca domestica*, is a particularly relevant insect to study in the context of the evolution of immune systems. House flies are versatile mechanical vectors of numerous diseases of human and livestock, including bacterial, protozoan, viral, and helminthic infections ranging from cholera to tapeworms (Scott et al. 2009; Joyner et al. 2013; Nayduch et al. 2013). Compared to other sequenced insects, they inhabit an unusually wide range of septic matter, including excreta, garbage, and diverse animal carcasses. This lifestyle suggests that house flies contact and must successfully avoid a wide range of potentially damaging bacteria (Gupta et al. 2012), suggesting that house flies may have an unusually effective immune system to cope with these challenges.

House flies are also an ideal system for studying the comparative genomics of insect immunity because of their phylogenetic position among Dipterans (Scott et al. 2009). The mosquito clade and the Drosophilids have been very heavily sampled for genome sequencing, but these two groups diverged approximately 250 million years ago (timetree.org) and represent close to the maximal divergence among Dipterans. House flies split this deep phylogenetic branch between Drosophilids and mosquitos, and thus provide significant additional resolution to Dipteran genomics.

In this study, we generated new RNA-seq data from experimentally infected and control (sterile-wounded) house flies. With these new data, we characterized the transcriptional response to infection in *M. domestica*. When combined with existing genomic resources in house flies and other Dipterans, these data reveal a striking expansion in the recognition and effector repertoires in *M. domestica*. We also develop a new statistical model for inference of gene family evolution, and show that these expanded repertoires in house flies are most likely associated with extremely elevated rates of gene duplication specifically in immune gene families along the house fly lineage, suggesting that the unusual lifestyle of house flies may be driving increased diversification of immunological molecules.

## Methods

### 1. Data collection

In order to detect genes induced by infection in *M. domestica*, we infected adult female flies 4 days post-eclosion with a 50:50 mixture (by volume of O.D. 1.0 samples) of *Serratia marcescens* and *Enterococcus faecalis.* These are same bacterial strains used in previous similar studies (Sackton et al.2013), and were chosen to capture both Gram-positive and Gram-negative responses. Bacteria were delivered by pricking the thorax with a 0.1 mm dissecting pin to penetrate the cuticle of the flies. Control flies were pricked using the same protocol, but with sterile LB broth instead of bacterial cultures. Both control and infected flies were infected between 12:00-1:00 PM in a single day and frozen in liquid nitrogen 6 hours after treatment in pools of 5 flies.

To inform our analysis of the transcriptional response to infection in *M. domestica*, we also generated RNA-seq data for infected and control *D. melanogaster*. For the *D. melanogaster* study, we used the same bacterial strains and concentration as above and did the experiments with females 3-5 days post-eclosion, but control flies remained unpricked and flies (control and infected) were frozen 12 hours after treatment.

Subsequently, we extracted RNA from whole frozen flies in TRIZOL following standard protocols. RNA-seq libraries were made using the Illumina TruSeq RNA sample prep kit, and sequenced on a HiSeq 2500 platform.

### 2. Updating *M. domestica* gene annotations

We first sought to update the existing *M. domestica* gene annotations to detect gene models that might have been missing in the initial published annotation. To do this, we used a pipeline based on the Trinity-assisted PASA workflow (Haas et al. 2003; Haas et al. 2011; Haas et al. 2013) described at http://pasa.sourceforge.net/ and in more detail at the Github page associated with this manuscript (https://github.com/tsackton/musca-immunity). We started with the *Musca domestica* GFF, protein, and transcript files produced by NCBI during the initial annotation of the *Musca* genome (NCBI release 100) (Scott et al. 2014), available at ftp://ftp.ncbi.nlm.nih.gov/genomes/Musca_domestica/ARCHIVE/ANNOTATION_RELEASE.100/.

After running the PASA pipeline (https://github.com/tsackton/musca-immunity/tree/master/supplemental_methods/pasa), our primary goal was to extract novel gene annotations: we only added new gene models that may have been excluded from prior annotation, and did not to update existing gene models. The rationale for this decision is that in the absence of paired-end data or higher coverage data, determining biologically real novel splice forms is a challenging problem subject to a high false-positive rate. Thus we focused exclusively on novel gene annotations, that is, gene models predicted by PASA from Trinity alignments to the *M. domestica* genome that do not overlap existing annotations. We identified 70 new protein-coding transcript models with this approach. Although by definition these tend to be predicted proteins with little homology evidence (as genes with strong homology to other Dipterans would likely be annotated by the NCBI pipeline), and they are significantly shorter than previously annotated proteins (median length 223 aa vs. 389 aa, P=1.58x10^-6^, Mann-Whitney U test). The transcripts encoding these novel predicted proteins tend to be more highly expressed than those encoding previously annotated proteins (adjusted count 186 vs. 98, P=0.00014, Mann-Whitney U test). An updated GFF file, isoform-to-gene key, protein fasta file, and transcript fasta file are available as supplemental data and online at https://github.com/tsackton/musca-immunity/tree/master/input_data/annotations

#### 3. Differential expression analysis

We used RSEM to quantify differential expression after infection in *M. domestica* and in *D. melanogaster.* Briefly, we first trimmed reads using Trimmomatic, then computed expression for each transcript in our updated annotation described above using RSEM v1.2.16 (Li and Dewey 2011) using bowtie2 as the read mapper. The full code to run our RSEM pipeline is available at https://github.com/tsackton/musca-immunity/tree/master/supplemental_methods/difexp, and the raw RSEM output is available in the supplemental data and at https://github.com/tsackton/musca-immunity/tree/master/input_data/rsem. To infer differential expression, we used DESeq2 (Love et al. 2014) with standard options. The full scripts for differential expression inference and related statistical analysis are available at https://github.com/tsackton/musca-immunity/tree/master/R.

### 4. Bioinformatic characterization of predicted *M. domestica* proteins

We focused on characterizing three properties of *M. domestica* proteins that can be determined from sequence and comparative information: the presence of a signal peptide, the phylogenetic age of the gene, and the presence of immune-related protein domains. All scripts are available at https://github.com/tsackton/musca-immunity/tree/master/supplemental_methods.

To identify signal peptides, we used signalp v4.1 with default options run on all predicted *M. domestica* proteins.

To define phylogenetic age (specifically, phylostratigraphic age, *sensu* (Domazet-Loso et al. 2007))), we started with a series of blastp searches and defined age as the node of the tree of life at which the most distant blastp hit is detectable. This is conservative in the sense that we do not screen for any kind of parsimonious pattern, so a spurious deep hit will mean we consider a protein to be ancient even in the absence of any more closely related hits. When we say a gene is young, we simply mean that no homologs can be detected by BLAST to older lineages; other factors, such as length or overall rate of sequence evolution, can thus impact gene age estimation if they increase the probability that distant homologs will be missed (Moyers and Zhang 2015). In particular, proteins that are rapidly evolving will tend to appear younger than their true age, and proteins that are short may also appear younger than their true age, due to biases inherent in detecting distant homologies of short and/or rapidly diverging sequences (Moyers and Zhang 2015). While in most cases our results focus on relatively recent homologs (*i.e* within Diptera or within insects), which are likely relatively unaffected by these biases (Moyers and Zhang 2015), we also corrected for these effects (at least partially) by modeling the impact of protein length and evolutionary rate (using expression level in *M. domestica* as a proxy) on our estimates of age. Formally, we first log-transformed and mean-recentered length and expression level, and then computed model coefficients for separate regressions with either scaled expression or scaled length as the predictor variable and age as the response. These coefficients are equivalent to the change in estimated age expected for a unit deviation from the mean (on a log scale) of either expression level or length. Length is essentially uncorrelated with estimated age in our dataset (Kendall’s tau = 0.02, P = 0.0002), but expression level is correlated with estimated age (Kendall’s tau =14 0.267, P < 2.2×10^-16^). To calculate scaled ages, we computed the normalized age as the real estimate age minus the predicted effect of expression; normalized age is no longer strongly correlated with expression, as expected (Kendall’s tau = -0.03, P =2.67×10^-9^).

To define phylogenetic age, we began with a curated set of complete proteomes (listed at https://github.com/tsackton/musca-immunity/blob/master/supplemental_methods/strata/strata_key.txt) and ran blastp against each complete proteome. For each set of BLAST results (representing the best hit of each *M. domestica* protein against a target database), we considered a hit as indicating the presence of a putative homolog if the alignment length is at least 40% of the *M. domestica* protein length and the alignment has at least 20% identity. We then extracted the deepest node for which we found evidence for a putative homolog, and defined that as the phylogenetic age of each *M. domestica* protein.

In order to quantify the presence of domains that have putative immune function, we first built a set of HMM profiles based on ImmunoDB curated alignments (http://cegg.unige.ch/Insecta/immunodb) (Waterhouse et al. 2007) and additional alignments for the Nimrod domain, IGSF proteins, and transferrins based on sequences downloaded from FlyBase. The non-ImmunoDB alignments, as well as the final alignment file of all immune-related proteins, is available in the Github repository associated with this paper. We then searched the complete set of predicted *M. domestica* proteins for matches to predicted immune-related HMMs using HMMER 3.0. We then processed the HMMER output to i) exclude cases where the E-value of the best domain is greater than 0.001, ii) the overall E-value is greater than 1x10^-5^, and iii) assign proteins that match multiple HMMs to the single HMM with the best e-value. To provide comparative information for the analysis of *M. domestica*, we also searched the predicted proteomes of the other Dipterans listed in Table S1 against our immune-related HMM database, and inferred the presence of domains with putative immune function using the same protocol.

### 5. Determining orthologs and paralogs of *M. domestica* proteins across Dipterans

To determine patterns of orthology and paralogy of *M. domestica* proteins among Dipterans, we built a gene-tree-based pipeline for identifying gene families and determining the relationships among genes. This pipeline is described in full at https://github.com/tsackton/musca-immunity/supplemental_methods/orthologyand in brief below.

First, we used OMA version 0.99 (Altenhoff et al. 2013) with default options to generate an initial set of homologous groups (HOGs), using as input the longest protein translation of each annotated protein in the *M. domestica* genome along with 13 other Dipterans (5 mosquitos, 7 Drosophilids, and *Glossina moristans*; Table S1).

After running OMA, we refined orthogroups as follows. First we generated an alignment of each initial orthogroup using MAFFT (Katoh and Standley 2013), and then created HMMs for each group using HMMER version 3. We then refined orthogroup assignment by searching each protein against each HMM, and merging orthogroups linked by a well-supported HMM hit. We also added genes to orthogroups when a gene was not initially assigned to any group, but has a significant HMM hit to a group (see part 1 of readme at Github site).

After orthogroup updating, we realigned each orthogroup with MAFFT (--auto option) and then computed a gene tree using RAxMLHPC-SSE3 version 7.75 (Stamatakis 2014), with the default options except -m PROTGAMMAAUTO and -N 10 (see part 2 of readme at Github site).

In some cases, our pipeline led to large gene families with one or more duplications at the base of Diptera. To both increase the computational efficiency of Treefix, and improve the accuracy of our rate estimation, we used a custom Perl script (treesplit.pl on Github) to split trees where the deepest node was inferred to be a duplication rather than a speciation event. After the first round of tree splitting, we used the programs Treefix (v. 1.1.8; default options except -m PROTGAMMAWAG, -niter=1000, and –maxtime) and tree-annotate (part of the treefix package) to reconcile the species tree with each gene tree and compute the likely number of gains and losses on the tree (Wu et al. 2012). Treefix attempts to produce the most parsimonious tree with respect to duplications and losses while remaining consistent with the maximum likelihood gene tree. It does this by searching the neighborhood of the maximum likelihood tree for topologies that reduce the number of duplication and loss events without significantly reducing the likelihood of the tree under the evolutionary model specified. Because this process is inefficient on large trees, we set a maximum time for the program to run (∼1 week), which means that for large trees we sample fewer iterations than for small trees. To partially account for this, we ran a second round of tree splitting with our treesplit script after our first round of TreeFix (which led to some large trees being split into smaller trees), and then repeated treefix on any altered trees. We then ran tree-annotate to produce duplication/loss inference on this final set of trees.

### 6. Analysis of gene family dynamics

To determine rates of gene duplication and loss across the phylogeny, we used both previously published, count-based methods such as CAFE (De Bie et al. 2006) and we implemented a Poisson regression model using duplication and loss events inferred from gene tree / species tree reconciliation. To account for differences in branch lengths, we constructed an ultrametric tree as follows (https://github.com/tsackton/musca-immunity/tree/master/supplemental_methods/ultrametric). First, we identified orthogroups with no duplications or losses across the phylogeny. Second, we concatenated the trimmed alignments of these orthogroups to produce a single Dipteran alignment for tree estimation. Finally, we used RAXML (version 7.7.5) with the -f e option (to estimate branch lengths on a fixed phylogeny) to estimate branch lengths from the known Dipteran phylogeny. Finally, we used the “chronos” function from the ape package in R to convert the tree to an ultrametric tree with arbitrary edge units.

To test for variation in rates of duplication and loss among different classes of genes along different lineages, we use a mixed model Poisson regression. Specifically, we fit a model which includes both fixed effects (functional class, lineage of interest), branch length as an offset, and a separate random intercept for each gene family, to control for overdispersion caused by rate variation among gene families, using the “glmer” function in the R package “lme4”. R code to implement this approach, and containing the full models used for each analysis, is available at https://github.com/tsackton/musca-immunity/tree/master/R. This approach allows us to use the full power of general linear models to test hypotheses concerning lineage-specific rates of duplication.

To test our Poisson regression approach, we simulated 1000 trees each with one of twelve different rates of gene duplication (assuming equal birth and death rates), ranging from 0.00057 events/MYA to 0.3412 events/MYA. To do these simulations, we fixed the species tree and estimate a gene tree within the species tree using the GuestTreeGen tool (part of jprime) with options -minper 0-min 4 -maxper 10000 -max 10000 (code: https://github.corn/tsackton/musca-immunity/tree/master/supplementalmethods/sims). The simulation approach of GuestTreeGen is based on a duplication-loss model where duplications and losses each occur with a specified Poisson rate along branches of a phylogeny, and speciation events result the simulated lineage splitting into two child lineages that continue to evolve by duplication and loss independently (Sjöstrand et al. 2013). After simulating data, we estimated gain/loss rates using both CAFE and our Poisson regression in order to estimate the duplication, or duplication and loss rates independently, for each simulated dataset. In order to calibrate the statistical properties of our regression approach, we also simulated 1000 data sets in which a random sample of 100 trees with different rates were selected to represent “immune genes.” We then test whether we find a significant difference between rates of duplication in “immune genes” compared to “non-immune genes”, using the Poisson regression approach described above.

## Results

### *Identifying genes regulated by infection in* Musca domestica

To characterize the infection-regulated transcriptome in *Musca domestica,* we used RNA-seq to quantify expression of genes and transcripts in infected and control (sterile-wounded) flies. We infected 4 day old adult female flies by piercing the cuticle with a dissecting pin dipped a mixed bacterial culture of *Serratia marsecens* and *Enterococcus fae calis.* Control flies were treated identically, except they were poked with a pin dipped in sterile LB broth. Six hours after treatment we collected three replicate pools each of infected and control flies, and sequenced each pool using standard Illumina protocols. Combined, we sequenced 45.5 million reads from infected flies and 51.0 million reads from control flies, of which roughly 70% map to *M domestica* gene models (NCBI annotation version 100) using RSEM (Li and Dewey 2011).

We identified genes differentially regulated between control and infected samples using the negative binomial approach implemented by DESeq2 (Love et al. 2014). We are able to detect expression for 13,621 genes, out of 14,466 annotated genes in the genome. Overall, we find 1675 genes differentially regulated at a 5% FDR, with 784 upregulated and 891 downregulated (Figure 1), representing 5.4% and 6.2% of genes in the genome, respectively.

**Figure 1.**
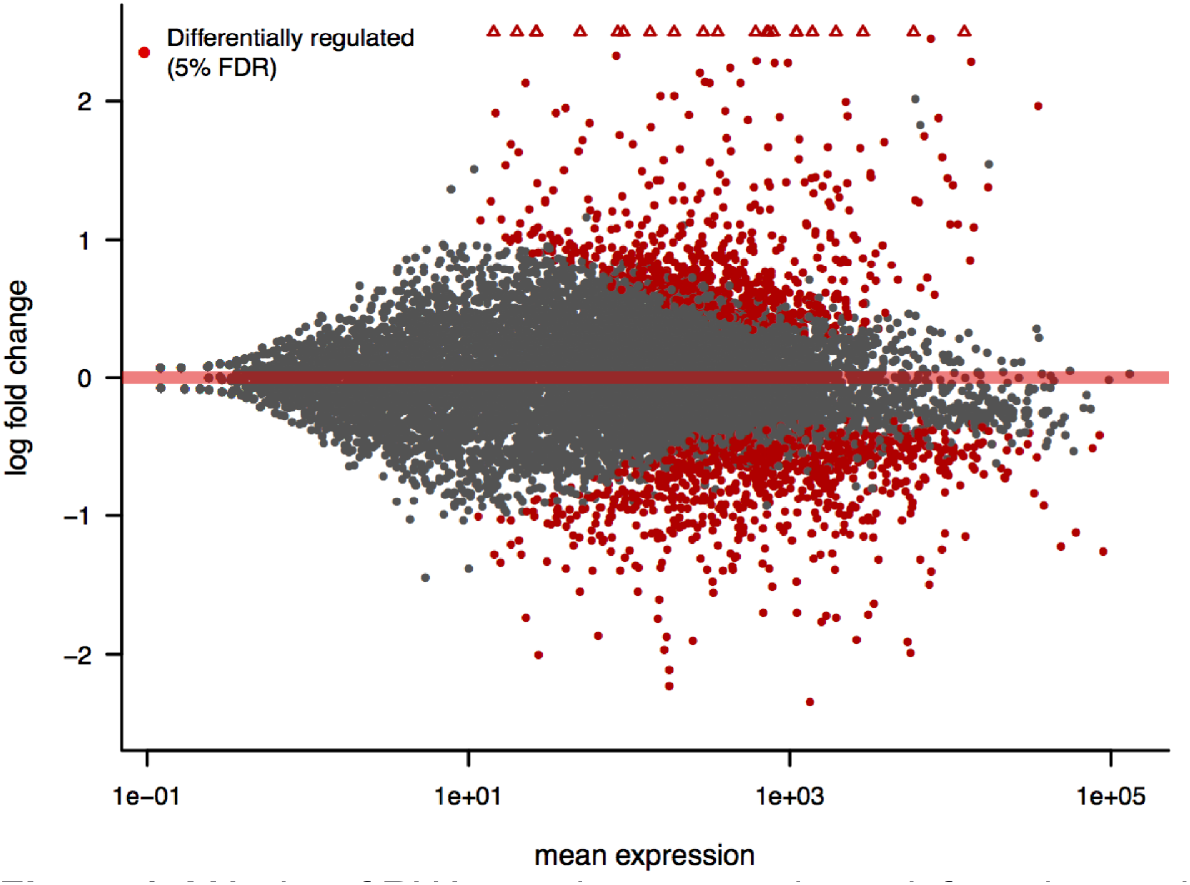
MA plot of RNA-seq data comparing uninfected control (sterile wounded) samples to infected samples. The x-axis shows mean expression for each M. domestica gene (as estimated in DESeq2), and the y-axis shows log2 fold change (infected vs. uninfected), also estimated in DESeq2. Points in red are differentially regulated between treatments at a 5% FDR. Open triangles represent points with log2 fold change greaterthan 2.5.

We used two approaches to identify genes in *M. domestica* with homology-based evidence for an immune function. First, we screened for homology to a curated list of genes with immune function in *D. melanogaster* (Table S2). Second, we used an HMM-based approach (Waterhouse et al. 2007) to identify house fly proteins with homology to previously characterized Dipteran immune-related gene families. The gene families we analyzed are listed in (Table 1), and alignments and HMMs are available online (https://github.com/tsackton/musca-immunity/tree/master/supplemental_methods/hmm). As expected, these homology-annotated immune genes have a much higher proportion of induced genes than the set of expressed genes as a whole (Dmel homology: 25.8% induced; HMM: 15.8% induced; all expressed genes: 7.6% induced, both comparisons P < 2.2×10^-16^, Fisher’s Exact Test).

**Table 1.**
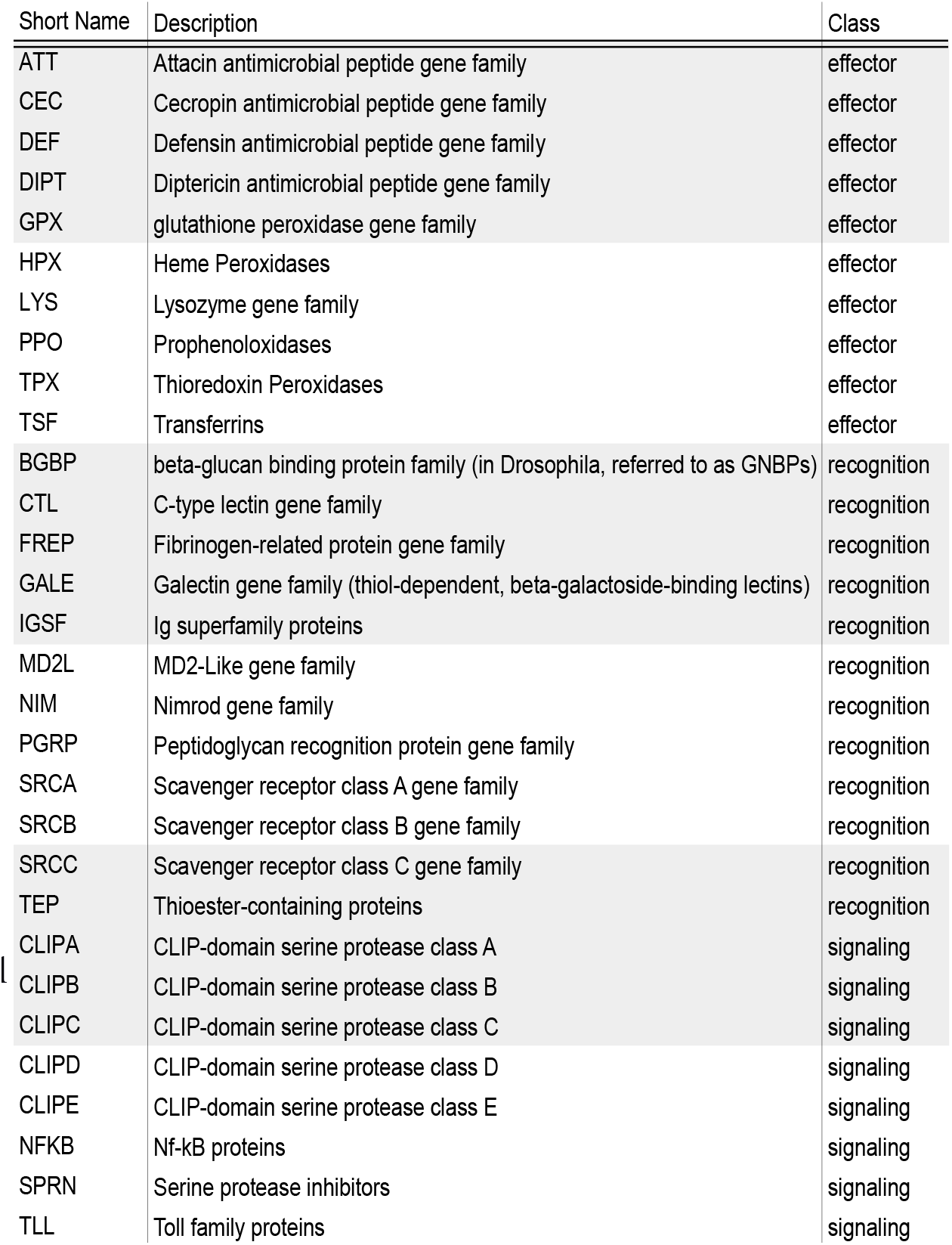
Immune-related gene families annotated by Hidden

Looking at individual genes induced by infection in *M. domestica* reveals a clear enrichment for genes with well-characterized immune annotations (Figure 2A). These include many homologs of consistently and strongly induced effector genes in *D. melanogaster*, such as cecropins (7 gene family members induced more than 2-fold in *M. domestica*), attacins (5 family members induced more than 2-fold in *M. domestica*), diptericins (2 family members induced more than 2-fold in *M. domestica*), and defensins (2 family members induced more than 2-fold in *M. domestica*). These also include homologs of genes, such as FREPs (8 induced in *M. domestica*) and galectins (5 induced in *M. domestica*) that have immune roles in some animals (Adema et al. 1997; Vasta 2009; Romero et al. 2011), including mosquitos (Dong and Dimopoulos 2009), but have not been experimentally characterized in *Drosophila*. A full list of genes with expression information is available at https://github.com/tsackton/musca-immunity/blob/master/results/mdom.difexp.tsv.

**Figure 2.**
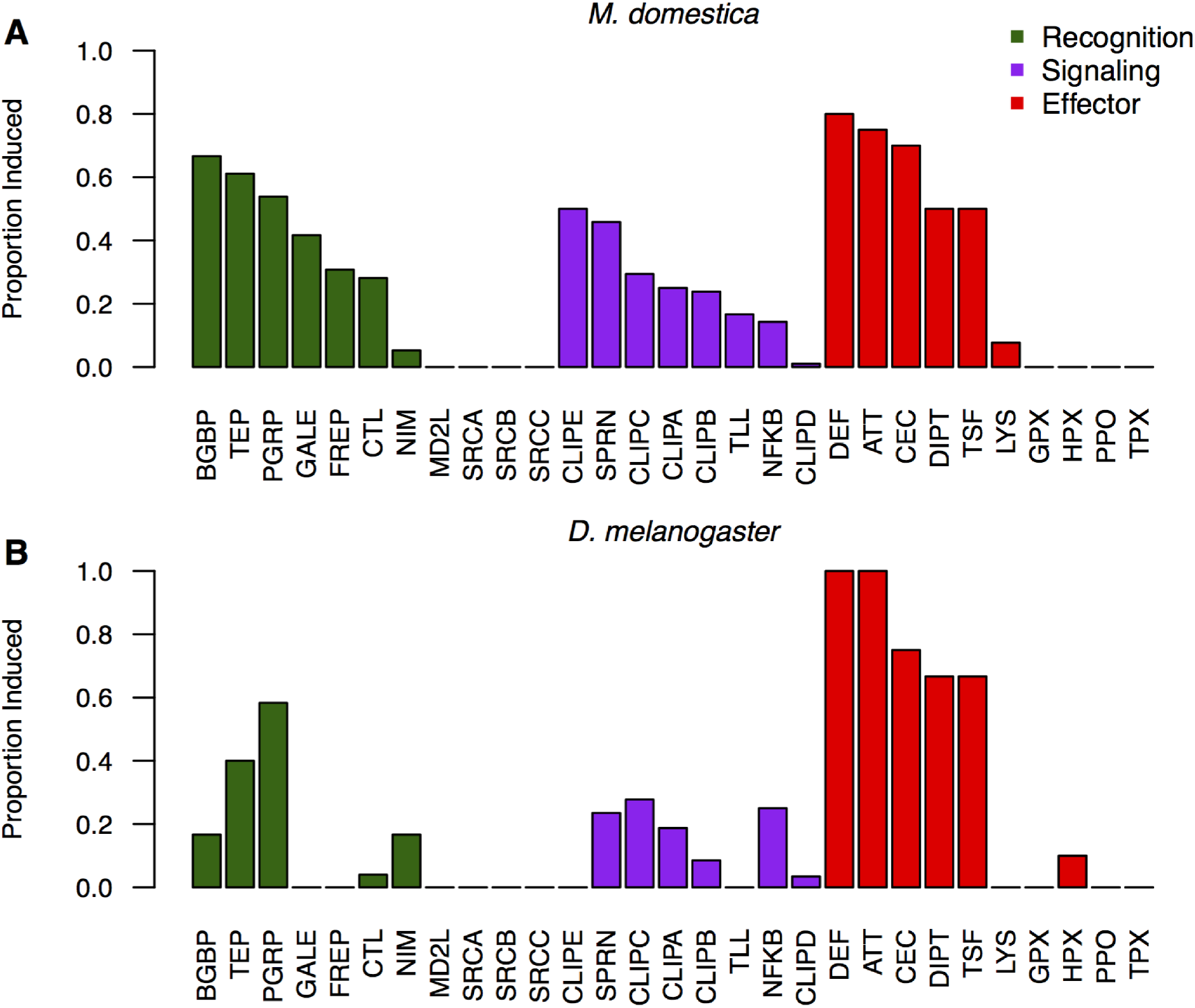
A) The proportion of each family of immune-related genes that are upregulated (at a 5% FDR, based on DESeq2 analysis) after infection relative to uninfected control (sterile wounded) at 6 hours post treatment in M. domestica. Families are defined based on HMM profiles and ordered by category (recognition, signaling, effector) and proportion induced within each category. B) The proportion of the each family of immune-related genes that are upregulated (at a 5% FDR, based on DESeq2 analysis) after infection relative to an uninfected control (naive, untreated) at 12 hours post treatment in D. melanogaster. Families are ordered as in part A.

Combining all sources of evidence (HMMs, *D. melanogaster* homology, gene ontology, and regulation after infection), we identify and annotate a total of 1,392 putative immune-related genes in *M. domestica.* A full list Markov Models. of these genes, with annotations where possible, is available as Table S3.

### Gene ontology analysis suggests a coordinated shift from metabolism to protein production after infection

In addition to genes encoding proteins with specific immune functions, bacterial infection leads to broad changes in patterns of gene expression that may be reflective of physiological processes altered by infection. To better understand the overall biology of the transcriptional response to infection in house flies, we focused on the 613 induced genes (FDR < 0.05) and 568 repressed genes (FDR < 0.05) which were able to be annotated to GO terms based on homology (Scott et al 2014).

As expected, genes induced by infection are enriched for GO classes related to immunity, including “response to biotic stimulus” (Holm’s adjusted P-value = 4.23×10^13^, Odds Ratio = 2.55), “response to stress” (adjP = 1.82×l0^-09^, odds ratio = 1.80), and “response to external stimulus” (adjP = 5.22x 10^-03^, odds ratio = 1.52). Additionally, genes induced by infection are enriched for a number of 8 biological process GO categories that are suggestive of a coordinated upregulation of protein synthesis and export machinery. These include “translation” (adjP = 4.98×10^-04^, odds ratio = 1.98), “transport” (adjP = 2.17×l0^-02^, odds ratio = 1.35), “cellular protein modification process” (adjP = 4.16×l0^-02^, odds ratio = 1.39), and “protein metabolic process” (adjP = 7.70×10^-07^, odds ratio = 1.63).

In contrast, genes repressed by infection are enriched for GO terms suggestive of a role in metabolism. GO terms overrepresented in the downregulated gene set are primarily related to 14 metabolism: “generation of precursor metabolites and energy” (adjP=1.32×l0^16^, odds ratio = 4.11), “lipid metabolic process” (adjP=5.79xl0^-09^, odds ratio = 2.13), “catabolic process” (adjP=3.24×l0^-08^, odds ratio = 1.83), and “secondary metabolic process” (adjP=6.06×10^-03^, odds ratio = 2.16). Molecular function GO terms paint a similar picture (Table S4). Taken together, these patterns point toward a pronounced physiological shift in house flies after infection, away from basal metabolism and toward protein production, transport, and secretion. This is consistent with recent work in *Drosophila* and other insects suggesting a close connection between metabolic control and immune system regulation (De Gregorio et al. 2001; Buchon et al. 2014; Unckless et al. 2015). A full list of GO terms enriched (at a Holms-adjusted P-value < 0.05) for genes either upregulated or downregulated by infection is in Table S4.

### *Comparison to* D. melanogaster *RNA-seq data suggests* M. domestica *induces a larger suite of genes after infection*

To contextualize our observations about the genes induced by infection in *M. domestica,* we generated in parallel a new, roughly comparable *D. melanogaster* RNA-seq dataset. While previous studies have been conducted of the transcriptional response to infection in *D. melanogaster* (De Gregorio et al. 2001; Irving et al. 2001), a direct comparison has the benefit of using data generated with the technology, the same infection protocol, at a similar time point, and in the same laboratory as the *M. domestica* data (see methods for details), minimizing technical artifacts. We also used the exact same analysis pipeline to analyze the *D. melanogaster* RNA-seq data. The RNA-seq data from *D. melanogaster* is of roughly similar depth and quality (67.8 million reads for the infected replicates pooled, 75.6 million reads for the uninfected replicates pooled, 95% mapped to *D. melanogaster* gene model); the only differences are 1) we sampled flies 12 hours after infection, instead of 6 hours, and 2) we used an untreated control instead of a sterile-wounded control. Both of these differences are likely to increase the number of genes detected as regulated by infection in *D. melanogaster*.

Of the 11,135 genes in *D. melanogaster* with detectable expression in our data, 156 are upregulated by infection and 150 are downregulated by infection, representing 1.4% and 1.35% respectively of expressed genes, and 0.9% and 0.87% respectively of all genes. This is notably fewer than in *M. domestica*, especially when taking into account the likely lower quality of the house fly annotations. Of induced genes, 27.6% are annotated as having an immune function. Unsurprisingly, the induced genes include many known antimicrobial peptides (4 attacins, 3 cecropins, defensin, 2 diptericins, drosomycin, and drosocin), recognition factors (2 Teps, 7 PGRPs, and 2 Nimrods), and signaling components (cactus, Relish). A full list of genes with expression information is at https://github.com/tsackton/musca-immunity/blob/master/results/dmel.difexp.tsv. At the level of HMM-defined gene families, *D. melanogaster* induces many of the expected classes, with substantial overlap with the classes induced in *M. domestica* (Figure 2B). Notably, however, we find no evidence for induction of any FREP or galectin in *D. melanogaster*, in contrast to the 31% and 42% respectively of genes in these classes induced by infection in *M. domestica*.

In our dataset there are 7,934 single-copy orthologs between *D. melanogaster* and *M. domestica* with detectable expression in both species. For these genes, we directly compared patterns of regulation after infection. While we find, as expected, highly significant overlaps in both induced genes (P=4.61×10^-12^, Fisher’s Exact Test) and repressed genes (P=4.1×10^-07^, Fisher’s Exact Test), there are many more genes induced in *M. domestica* alone than in *D. melanogaster* alone (Figure 3). This suggests that at least a portion of the greater number of genes regulated by infection in *M. domestica* is attributable to regulatory evolutionary change in shared orthologs.

**Figure 3.**
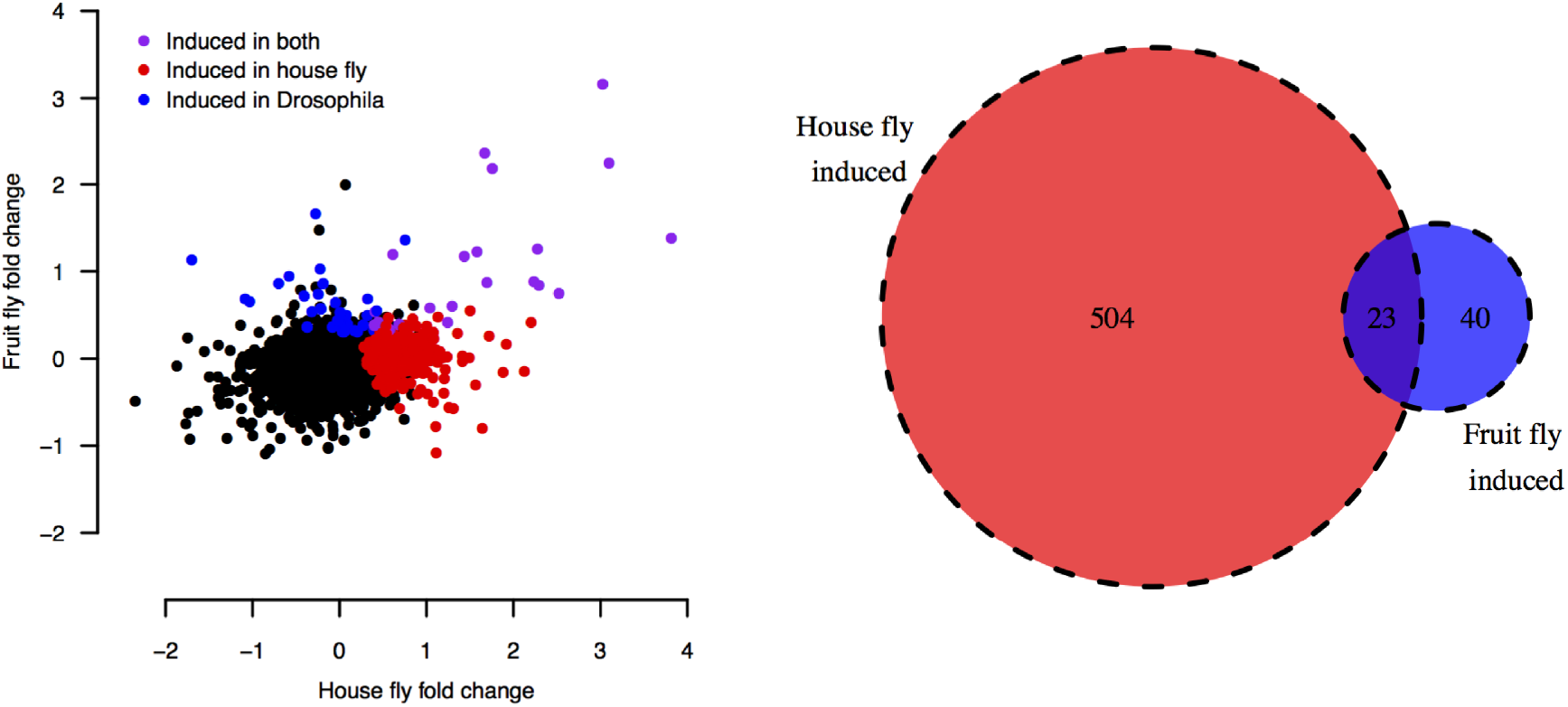
The correlation between fold change after infection in house fly and fruit fly. Each point represents a single 1:1 ortholog with expression data in both species. Genes with significant upregulation after infection in either fruit fly, house fly, or both are colored; significant upregulation is defined based on a 5% FDR estimated with DESeq2. The Venn diagram shows the number of 1:1 orthologs in each induction class.

We also compared the set of gene ontology terms overrepresented among both upregulated and downregulated genes in *D. melanogaster* to those described for *M. domestica* above. In *D. melanogaster*, GO terms associated with immune functions dominate the list of terms overrepresented in the upregulated class (Table S4). However, we see no evidence for upregulation of GO terms associated with protein transport or translation. It is possible that differences in timing (6 hours vs 12 hours) could be associated with this difference, but it is also possible that this represents a reduced investment in immune protein production in *D. melanogaster* compared to *M domestica*. For the downregulated genes, we see a similar set of GO categories associated with the *D. melanogaster* response as the *M domestica* response (Table S4), supporting the idea that the downregulation of basal metabolism is a broadly consistent response to infection in many Dipterans.

Taken as a whole, bacterial infection in *M domestica* appears to result in differential expression of more genes that in *D. melanogaster*(including single-copy orthologs that are not regulated by similar bacterial infections in *D. melanogaster).* These additional regulated genes appear to include additional categories of immune-related genes (FREPs, galectins), a broader range of biological processes (including protein translation and export machinery), and induction of more members of shared immune related families that may have expanded in *M. domestica* (including attacins, cecropins, TEPs, transferrins, defensins).

### *The infection-induced transcriptome of* Musca domestica *is enriched for taxonomically young genes*

In several insects studied to date, the transcriptional response to infection includes a large number of young, taxonomically restricted genes (Sackton and Clark 2009; Sackton et al. 2013; Gupta et al. 2015). To test whether the data from *Musca domestica* also show this pattern, we identified the phylogenetic age of each protein in the house fly genome using BLASTP and then inferring a date for gene origination based on the age of the deepest homolog identified (ages from timetree.org). As has been seen in other insects, young genes in *Musca domestica* are more likely to be induced by infection than old genes (Logistic regression: β= -5.53×10^-4^ P=7.88×10^-13^Figure 4A).

**Figure 4.**
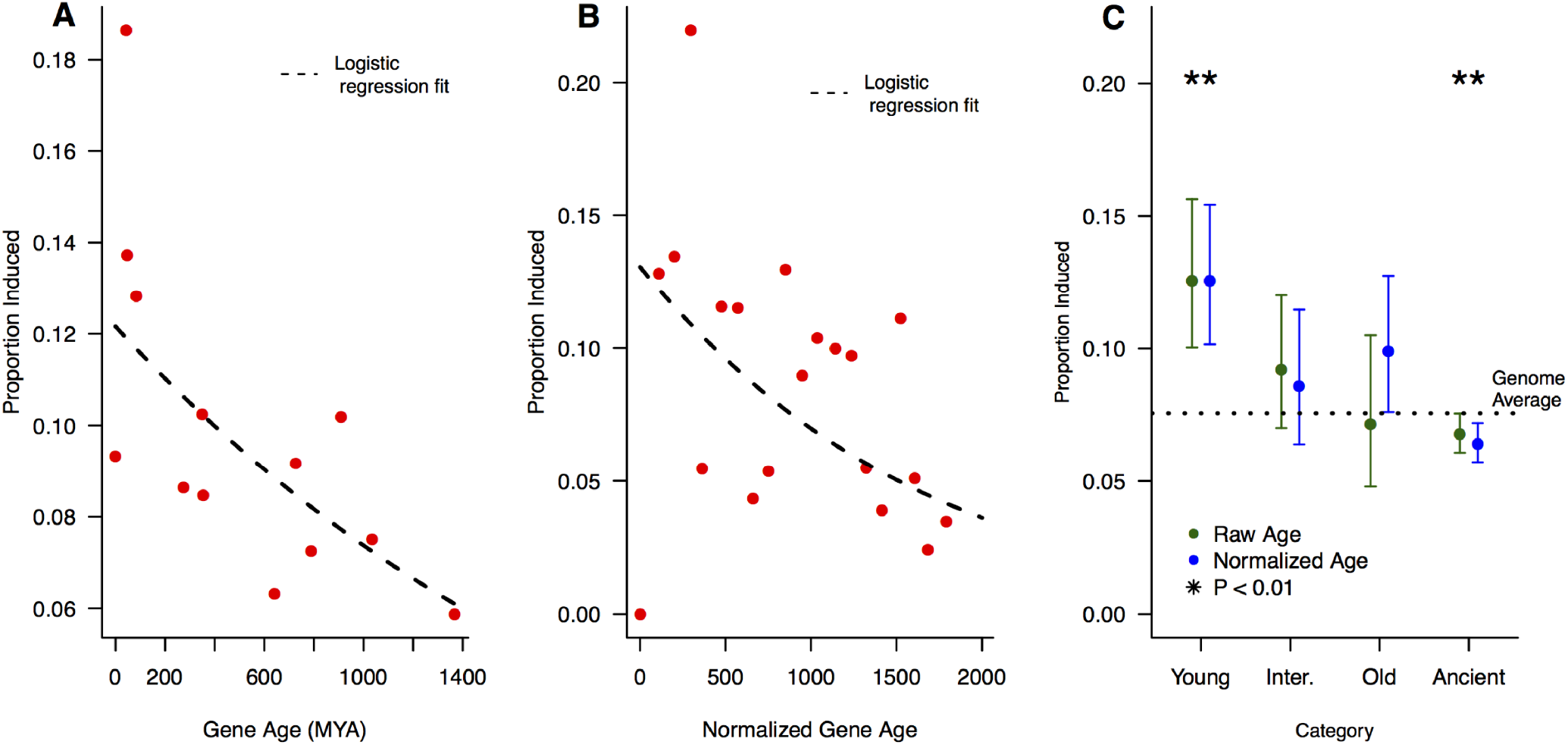
A) The proportion of genes induced by infection for each inferred gene age. The dashed line shows the logistic regression fit, which is highly significant (age β= -5.53×10^-04^, P = 7.88×10^-13^). B) The proportion of genes induced by infection for each normalized gene age. The dashed line shows the logistic regression fit, which is highly significant (age β = -6.94×10^-04^, P = 2×10^-16^). Note that the normalization procedure generates a continuous distribution of ages, but for plotting purposes we converted this back to discrete age classes. C) Proportion of genes induced by infection by age category. After classifying genes into one of four categories based on either raw (uncorrected) age (green points) or normalized (corrected) age (blue points), we estimated the proportion of each age class induced by infection. The dotted line shows the genome-wide average proportion genes induced by infection (0.081). To estimate significance, each category was compared to the remaining categories in turn using a chi-square test. We get similar results using a logistic regression to estimate the effect of each category relative to the “ancient” group.

Recently, it has been suggested that phylostratigraphic methods such as this are prone to bias, since factors such as protein length and evolutionary rate can influence the probability of detecting ancient homologs (Moyers and Zhang 2015). To attempt to control for this effect, we normalized our age estimates based on the estimated effects of protein length and expression level as a proxy for evolutionary rate (Pál et al. 2001; Larracuente et al. 2008) in our data, and repeated our analysis (see methods for details). After this correction, we still find strong evidence that younger genes are more likely to be induced by infection than older genes (Logistic regression: β = -6.93×10^-04^, P < 2×l0^-16^,Figure 4B).

As an alternative approach, we also assigned genes to a small number of age categories (young= Schizophora-specific genes, intermediate = Insecta-specific genes, old = Protostomia-specific genes, ancient = Opisthokonot-specific genes) and consider the patterns of expression in genes in each category. Using both uncorrected and corrected phylostratigraphic age categories, we find that genes in the ’young’ or ‘intermediate’ categories are more likely to be induced after infection than genes in the ‘old’ or ‘ancient’ categories (Figure 4C).

### *Gene duplication and loss in* Musca domestica

In addition to apparently inducing a broader suite of genes encoding immune-related proteins than many other insects, the *Musca domestica* genome encodes a greater diversity of immune-related genes than many other insects studied to date. For example, the *Musca* genome contains the highest number of TEPs in a sequenced Dipteran genome (Scott et al. 2014), and in general has a high number of many immune-related gene families (Table 2). To test whether this is a general pattern across the *Musca* immune system, and to determine whether the diversity of immune proteins in *Musca* is driven by increased rates of gene duplication, decreased rates of gene loss, or both, we developed a phylogenetic framework to assess rates of copy number change using a Poisson regression approach (Koerich et al. 2008).

**Table 2.**
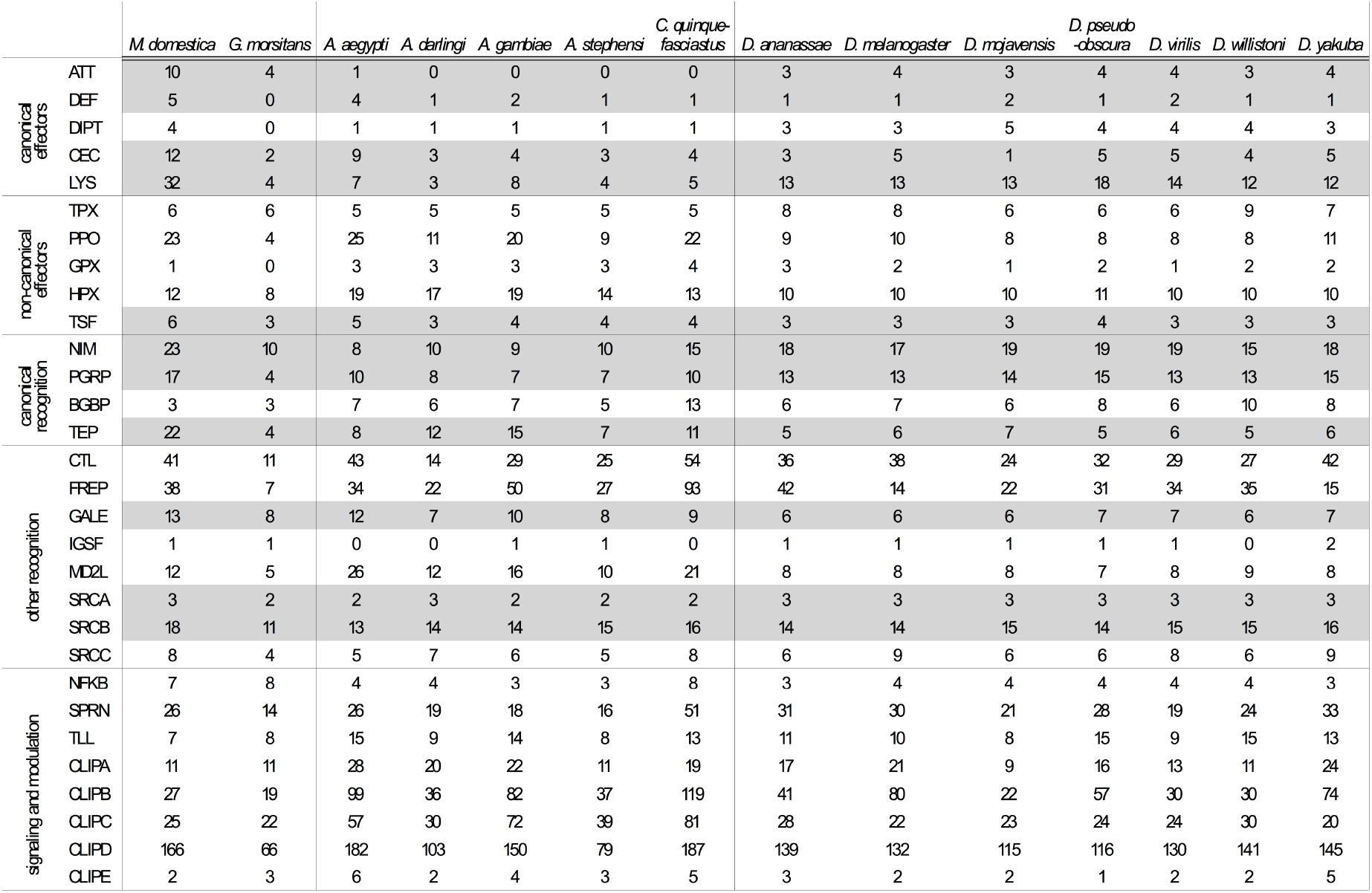
Number of genes identified by HMM for each gene family from Table 1. Rows where M. domestica has the highest count are shaded in gray.

In this framework, we fit a Poisson regression to counts of gene gains and gene losses on each branch of the Dipteran phylogeny (Figure 5). We first verified the behavior of our model by simulation and by comparison to previous methods. Then, we focused on three different model parameterizations. In the first approach, we allowed a different rate of duplication and loss on the *Musca* lineage compared to the rest of the tree, and estimated a single birth rate and death rate for all genes with a similar functional annotation (*e.g* recognition, signaling, effector, non-immune). In the second approach, we focus on specific related gene classes (e.g Cecropins, TEPs). Finally, we fitted a separate birth and death rate for each individual gene family. In all analyses, we focused on gene families basally present in Diptera, and with at least one gain or loss on the tree.

**Figure 5.**
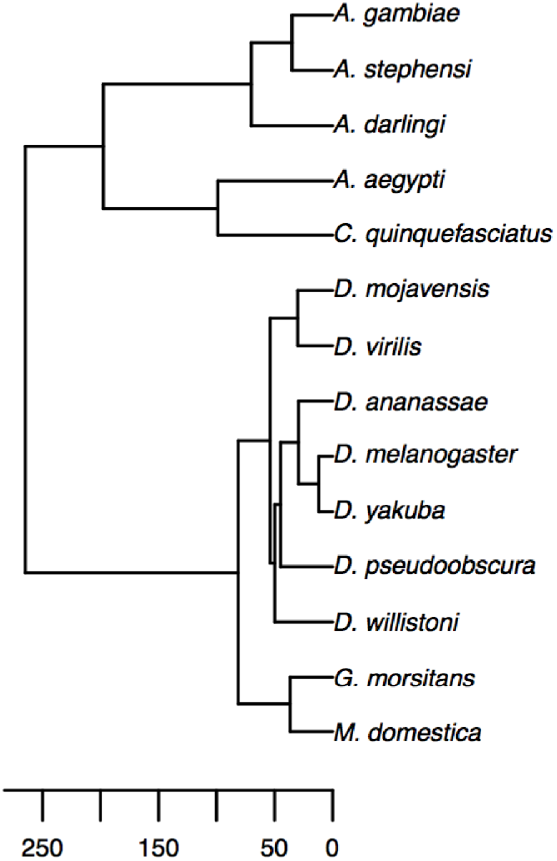
Ultrametric tree of Dipteran species included in gene family analysis, estimated using the “chronos” function in the ape package for R. Scale bar is in millions of years ago,based on the callibration taken from timetree.org.

### Poisson regression is an accurate method for estimating rates of gene gain and loss

To verify the behavior of our method, we simulated 1000 based on calibrations taken gene trees, conditioned on a fixed species tree, for each of 12 different duplication/loss rates ranging from 0.00057 to 0.341 (Table S5), using the GuestTreeGen tool injprime (Anon). In our simulations, we fixed the duplication rate to equal the loss rate (so the total rate in events / MY is twice the input simulation rate), and after simulation restricted our analysis to the subset of simulations where the gene family was not lost entirely on one of the two branches leading from the root of the tree (to be consistent with our filtering of our analysis of the real data).This drastically reduces the number of simulation results we used for the highest turnover rates (Table S5), but up to a turnover rate of 0.023 events / million years we retained at least 100 simulated trees. While we report results for all simulation values, those greater than 0.023 events / MY should be treated with caution due to the low numbers of gene families passing our filters.

For each set of trees simulated under the same rate parameters, we estimated a fixed turnover parameter (birth rate + death rate), and also separate birth and death parameters, using our Poisson regression model. Even for very high turnover rates, we recovered overall turnover rates and duplication rates very similar to the simulated values (Figure 6A). For low to moderate turnover rates, our estimates of loss rates were also very accurate, but for very high turnover rates we began to underestimate loss rates (Figure 6A), probably because losses that extinguish the gene family are dropped from the analysis and thus not counted. We note that existing methods such as CAFE also perform poorly at very high turnover rates (Figure 6B), and this is not unexpected (De Bie et al. 2006). At low to moderate turnover rates, our method performs as well as CAFE and allows for more complex modeling of branch dependencies.

**Figure 6.**
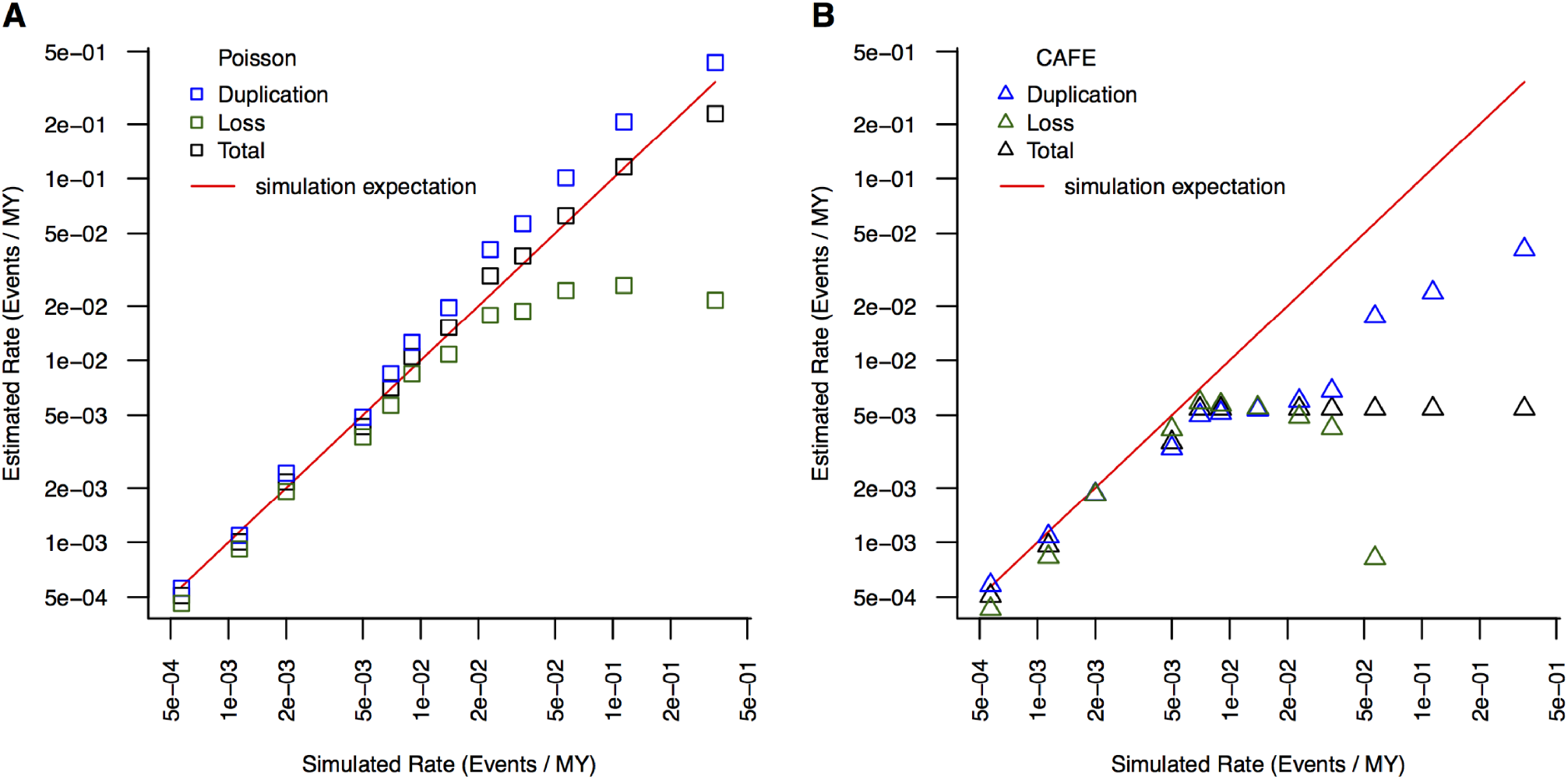
Analysis of simulated gene duplication data. The same simulation inputs (Table 85) were analyzed with our Poisson regression framework (A) and with CAFE (B) for a range of simulated duplication/loss rates. Points represent estimated turnover (duplication+ loss, black), loss (green) and duplication (blue) rates estimated with each method; the red line is the expectation based on the simulated input values.

In real data, considerable rate variation among individual gene families creates over-dispersion in the Poisson model and leads to serious underestimates of the standard error of the coefficients of the model, and thus incorrect P-values. To correct for this, we use a mixed model approach, specifying a random intercept for each individual gene family; this is conceptually similar to using observation-level random effects (Harrison 2014). To verify the performance of our mixed model, we simulated 1000 datasets with randomly selected “immune” genes, as described in the methods. On average the effect of this "immune" classification on duplication or loss rates should be zero in these random permutations, so we expect to observe no more than 5% of simulations that reject the null hypothesis of no effect at a nominal alpha of0.05. With the naive Poisson approach, we see a dramatic mis-calibration of the significance level (88.9% of all simulations have a P-value < 0.05), which is completely eliminated by accounting for family-level rate variation using random effects (5.1% of all simulations have a P-value 6 < 0.05).

### *Genes encoding effector and recognition proteins duplicate rapidly on the* Musca *lineage*

At the broadest level, we find evidence that the *Musca* lineage has experienced a significantly higher rate of gene turnover (duplication+ loss) than other Dipteran lineages for both immune genes (defined based on homology to *D. melanogaster*) and non-immune genes (Table 3). Notably, the increased turnover rate along the *Musca* lineage is significantly higher for immune genes than for non-immune genes (interaction β = 0.22, P = 0.0164), suggesting that immune genes in particular experience rapid turnover along the *Musca* lineage. The Musca-specific increase in turnover of in immune genes appears to be driven by an increased duplication rate (duplications only, interaction β=16 0.43, P =3.45×10^-05^) rather than a change in the rate of gene loss (losses only, interaction β= -0.0054, P= 0.983). When we define immune genes more broadly to include both genes with homology to *D. melanoga ster* immune-related genes and members ofHMM-defined immune gene classes, the same trends hold albeit somewhat more weakly (Table 3; duplications only interaction β = 0.31, P = 1.6×10^-04^ ; losses only interaction β = -0.26, P = 0.256).

**Table 3.** Poisson regression models for the analysis of duplication/loss data.

|  | Estimate | Std Error | z | Pr(>\|z\|) |
| --- | --- | --- | --- | --- |
| (Intercept) | -5.912 | 0.017 | -353.900 | <2e-16 |
| Lineage = Musca | 0.253 | 0.025 | 10.200 | <2e-16 |
| Immune (Narrow) = TRUE | 0.517 | 0.103 | 5.000 | 4.66E-007 |
| Lineage by Immune interaction | 0.222 | 0.093 | 2.400 | 0.0164 |

|  | Estimate | Std Error | z | Pr(>\|z\|) |
| --- | --- | --- | --- | --- |
| (Intercept) | -5.917 | 0.017 | -353.100 | <2e-16 |
| Lineage = Musca | 0.253 | 0.025 | 10.000 | <2e-16 |
| Immune (Broad) = TRUE | 0.529 | 0.086 | 6.200 | 7.07E-010 |
| Lineage by Immune interaction | 0.130 | 0.076 | 1.700 | 0.0868 |

To rule out the possibility that our results are driven by unusual rates in non-Musca lineages, we repeated these analyses with a model that allows for separate rates for each family of Dipterans included in our analysis (Muscidae, Drosophilidae, Glossinidae, and Culicidae), excluding events that occurred in basal lineages that pre-date the divergence of these families. In this analysis, we treated the Muscidae as the reference level; while genes with immune annotation have higher duplication rates in general than other gene families in the genome, in all non-Muscidae lineages the increase in duplication rates associated with immune function is significantly lower than the increase in duplication rates associated with immune function in Muscidae (family x immune interaction β = -0.58 for Drosophilidae, -1.136 for Glossinidae, and -0.62 for Culicidae, all P-values < 1× 10^-05^).

To initially determine if particular components of the innate immune system are responsible for this pattern, we estimated separate rates for different functional classes of immune proteins (recognition, signaling, modulation, and effectors; based on homology to *D. melanogaster* proteins with annotated functions in these classes). Gene families encoding recognition and effector proteins have elevated duplication rates in the Muscidae lineage compared to other Dipterans, but gene families encoding signaling or modulation proteins do not (Figure 7). Using simultaneous tests of linear contrasts, we tested whether this increase in immune-gene duplication rates is associated with increases in duplication rates in effector and signaling genes in models that allow separate rates for each major Dipteran lineage included in our dataset. We found the increase in duplication rates of genes encoding effector or recognition proteins (compared to the duplication rate of non-immune genes) is consistently elevated in the Muscidae lineage in all comparisons (Figure 7).

**Figure 7.**
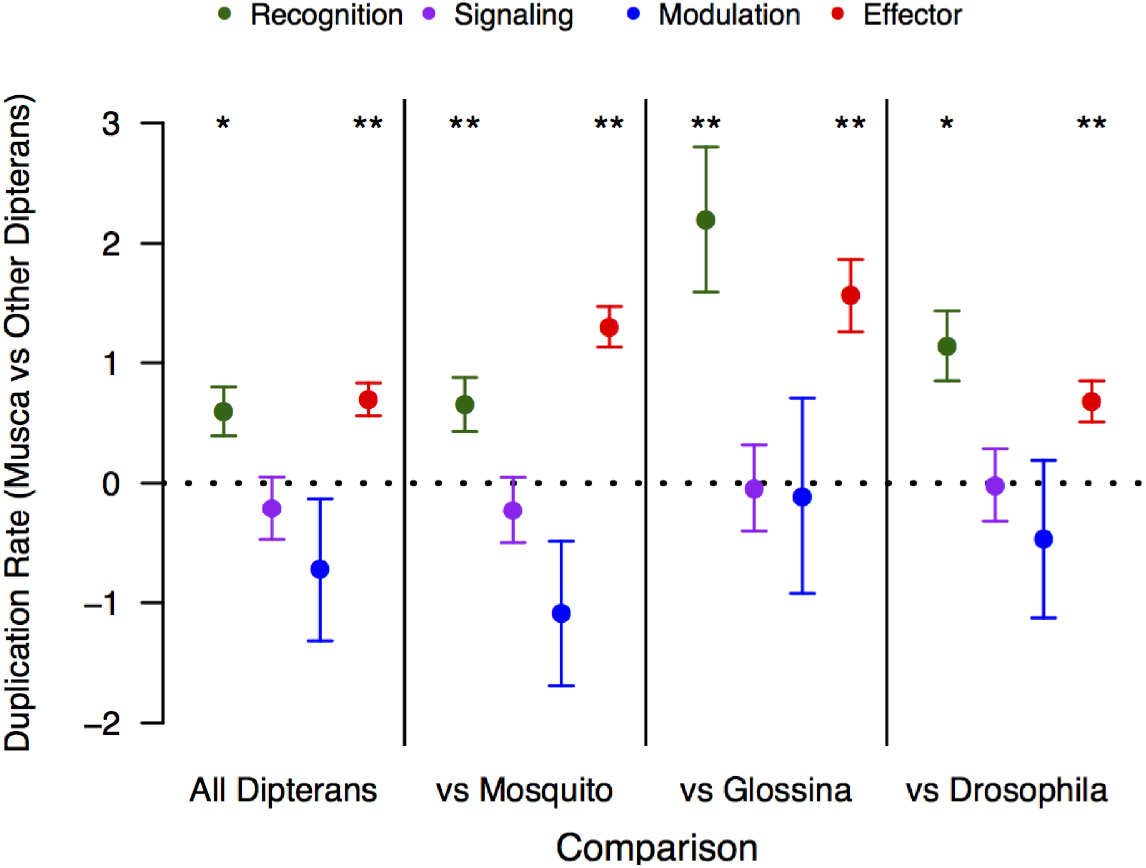
Linear contrasts testing the relative duplication rate in Musca lineage vs. other Dipterans for specific immune classes. Each point represents the estimated linear contrast(+/- standard error) for the duplication rate of genes in that category in Musca, compared to the duplication rate of genes in that category in all other Dipterans together or in individual non-Musca lineages. P-values are listed above each point for the testof whether the contrast is equal to 0, which is the expectation ifthe duplication rate for that category is equal on the Musca branch and the rest of the tree (* 0.01 < P < 0.05, ** P < 0.01).

In order to understand the specific drivers of this pattern, we analyzed rates of gene duplication and loss in HMM-defined gene families that make up the broader homology-based classes, focusing on effector and recognition classes. Among genes encoding recognition or effector proteins, the TEPs, lysozymes, and cecropins show the most striking pattern, with significantly larger increases in duplication rates (relative to the baseline non-immune duplication rate) along theM *domestica* lineage than in other Dipterans pooled (Table 4). Furthermore, for all three of these gene classes the increase in duplication rates in the Muscidae lineage is significantly greater than the increase in duplication rates in either the mosquito or the Drosophila lineages, relative to the baseline rate of all genes not in the family in question (Table 5).

**Table 4.** Linear contrasts testing duplication rate in the Musca lineage vs. other Dipterans for specific immune HMM families.

| Family | Estimate | Standard Error | P-value |
| --- | --- | --- | --- |
| BGBP | -4.034 | 8.026 | 1 |
| CEC | 1.794 | 0.4094 | 0.000294 |
| CLIPA | -3.185 | 8.95 | 1 |
| CLIPB | 0.06555 | 0.2382 | 1 |
| CLIPC | -3.608 | 10.76 | 1 |
| CLIPD | -0.7448 | 0.5169 | 0.982569 |
| CTL | 0.2163 | 0.2112 | 0.999891 |
| FREP | 0.1823 | 0.2149 | 0.999997 |
| GALE | -7.821 | 57.48 | 1 |
| HPX | -0.631 | 1.047 | 1 |
| IGSF | -0.101 | 1.051 | 1 |
| LYS | 1.427 | 0.2484 | <0.000001 |
| MD2L | 0.7331 | 0.3725 | 0.715598 |
| NFKB | 1.795 | 0.9337 | 0.754346 |
| NIM | -4.268 | 7.424 | 1 |
| PGRP | 1.201 | 0.4439 | 0.157238 |
| PPO | 1.036 | 0.3644 | 0.105604 |
| SPRN | 0.4289 | 0.3856 | 0.99956 |
| SRCA | -2.236 | 9.928 | 1 |
| SRCB | 0.1199 | 0.7669 | 1 |
| SRCC | 1.791 | 0.9597 | 0.798468 |
| TEP | 1.568 | 0.3018 | 0.0000515 |
| TLL | -4.964 | 8.636 | 1 |
| TPX | -79.75 | 47450000 | 1 |
| TSF | 2.887 | 1.245 | 0.402159 |

**Table 5.** Linear contrasts testing elevated duplication rate in the Musca lineage vs. other Dipteran families for cecropins, TEPs, and lysozymes.

| Family | Estimate | Musca vs Drosophila |  | Estimate | Musca vs Mosquitos |  |
| --- | --- | --- | --- | --- | --- | --- |
|  |  | Standard Error | P-value |  | Standard Error | P-value |
| TEP | 2.4461 | 0.6272 | 0.000192 | 1.2052 | 0.3312 | 0.000546 |
| CEC | 1.217 | 0.5174 | 0.036014 | 2.059 | 0.5486 | 3.48E-004 |
| LYS | 0.7971 | 0.2968 | 0.0143 | 2.9433 | 0.4924 | 4.52E-009 |

We can also fit our birth/death model to individual gene families (orthogroups), although in these cases we have substantially reduced power to estimate rates accurately, and thus will likely only detect the most extreme effects. We used this approach to estimate for each gene family the relative turnover rate (birth+death) on the Musca lineage compared to the rest of the tree; this is positive for gene families with a higher turnover rate on the Musca lineage and negative for gene families with a lower turnover rate on the Musca lineage. Immune-related genes (combining HMM-based and homology-to-Drosophila based annotations) are overrepresented among gene families with individually significant accelerations in turnover rate along the *M. domestica* lineage (6/154 immune families, 53/4565 non-immune families, P=0.012, Fisher’s Exact Test), including orthogroups containing TEPs, lysozymes, and cecropins (consistent with our HMM-class rate estimation; Table 6 has the full set of immune-related gene families with elevated turnover rates in Musca). Thus, all our modeling approaches consistently demonstrate a specific acceleration of rates of gene duplication in certain key classes of genes encoding recognition and effector proteins along the *M. domestica* lineage.

**Table 6.**
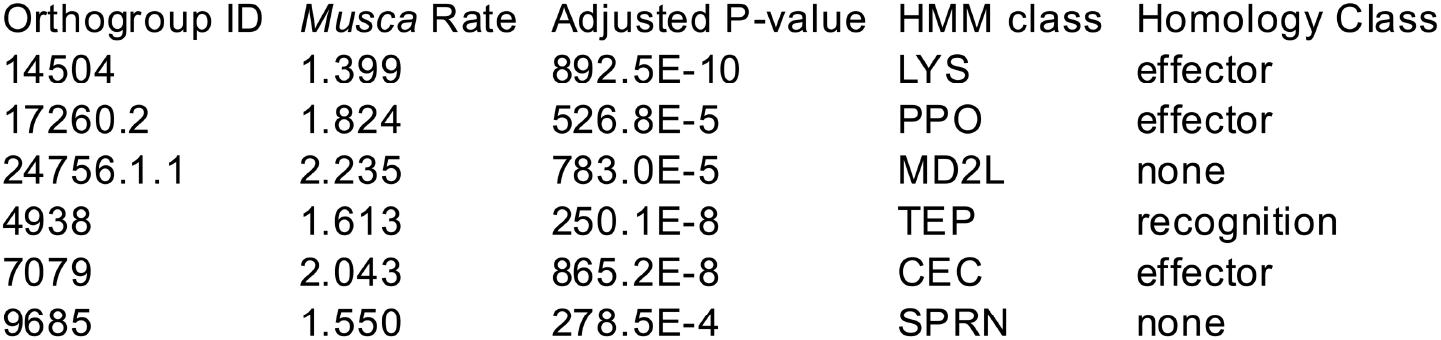
Orthologous groups with putative immune function and number of genes with evidence for accelerated duplication rates in the M. domestica lineage

As an additional line of evidence, we also examined the counts of each HMM-defined gene family detected in each species. Here, we don’t focus on rates of duplication or the phylogenetic relationships among genes, but rather just the absolute count of the number of genes with evidence for protein homology to particular immune gene families. We did this in as unbiased a way as possible, by using the same input set of HMM profiles to screen the full set of annotated proteins for each target species with HMMER. The counts of each gene family for each species are listed in Table 2. For all three gene families (Cecropins, TEPs, Lysozymes) where we infer a dramatically increased rate of gene duplication on the Musca lineage, we note that *M. domestica* has the most members of the full set of annotated Dipteran genomes we investigated. In general, the house fly immune system appears to encode a larger number of both effectors (including antimicrobial peptides, PPO pathway genes, and lysozymes) and recognition proteins (including TEPs, Nimrods, and PGRPs) than any other Dipteran included in our analysis.

## Discussion

The house fly, unique among Dipteran insects sequenced to date, lives primarily in highly septic environments, such as excreta, garbage, and carcasses. These environments have the potential to significantly impact the evolutionary dynamics of innate immune defense in this species. Organisms might deal with a potentially infectious environment by strengthening the barriers to initial infection, generating a more impermeable cuticle that is tougher or less prone to breaches that may allow bacterial invasion. They might also simply become more tolerant of bacterial presence, and not expend the energy entirely on strengthened resistance. In this study, we combine transcriptome sequencing before and after infectious challenge with homology based annotations to characterize the genes involved in the *M domestica* immune response and elucidate their evolutionary history. Numerous studies have reported that genes encoding proteins in the insect immune response are exceptionally likely to evolve by repeated positive selection (Schlenke and Begun 2003; Sackton et al. 2007; Lazzaro 2008; Obbard et al. 2009; Keebaugh and Schlenke 2012; Roux et al. 2014); here, we focus particularly on rates of gene gain and loss.

Several lines of evidence suggest that the *M domestica* immune response is unusual, at least when compared to the standard Dipteran model *D. melanogaster.* First, house flies appear to induce a broader range of putative immune genes than *D. melanogaster.* In addition to upregulating a number of conserved antimicrobial peptides (e.g defensins, cecropins) after infection, *M domestica* also induces large numbers of FREPs and galectins that are not induced in *D. melanogaster,* at least under the conditions we assayed. Second, we find some suggestion that house flies may induce a more robust immune response than *D. melanogaster* based on the function of non-immune genes that are regulated by infection. After challenge, we find that *M domestzca* upregulates a large number of genes with functions related to protein transport, protein synthesis, and 7079 2.043 865.2E-8 CEC effector protein export, and downregulates a large functions related to oxidative phosphorylation and metabolism. This pattern is consistent with a pronounced physiological shift of resources from basal metabolism to effector protein production and secretion. While the downregulation of genes with functions related to metabolism is likely a general response to infection across Dipterans, to our knowledge the upregulation of protein transport machinery has not be previously shown in Dipterans and is not detectable in our *D. melanogaster* expression data. In the absence of additional transcriptional data using the same challenges and experimental protocols in other Dipterans, it is of course formally possible that *D. melanogaster* is the atypical species in terms of 40 the transcriptional response to infection. However, our genomic analysis of gene duplication rates points to *M domestica* as the outlier.

Finally, we find clear evidence that genes encoding both recognition and effector components of the insect immune response are duplicating more rapidly along the *M domestica* lineage than in other Dipterans, and that among the gene families in its genome, those involved in recognition and effector functions are among the fastest to expand. This could be due to either selective or mutational processes, which are difficult to disentangle. It is tempting to speculate that this is driven by selection for either increased diversity or increased dosage in house flies, perhaps in response to their septic habitats. In an intriguing parallel, a high diversity of novel putative effectors is induced by LPS stimulation in the rat-tailed maggot (Altincicek and Vilcinskas 2007), which also inhabits a highly septic environment. Ultimately, however, more studies will be needed to test whether immune gene duplication rates are indeed increased generally in insects that live in particularly septic habitats.

More broadly, this study confirms the pattern observed in other insects that genes induced by infection have a general tendency to be taxonomically-restricted. However, what drives this pattern is still an open question. At least two hypotheses seem viable. First, it could be the case that young genes are in general less tightly regulated at the transcriptional level. As a consequence, in conditions of strong transcriptional activation (such as during an immune response), these genes have a tendency to be upregulated even without a clear function. Alternatively, this pattern could be driven by selective recruitment of novel genes to the immune system in response to the particular challenges that diverse insect lineages experience.

Ultimately, these conclusions solidify emerging evidence that rapid host-pathogen evolutionary dynamics are not limited to rapid sequence evolution. While it is difficult to know the ultimate cause of evolutionary change, this and other recent work makes clear that insect immune systems are extremely labile, not just at the level of protein sequence, but at the expression level and even at the level of gene content. It seems likely that much of these rapid changes are indeed driven by host-pathogen conflict, and that the evolutionary consequences of these arms races are broader than traditionally assumed.

## Acknowledgments

We thank Dr. Jeffrey Scott for providing the flies and husbandry expertise, Dr. Angela Early for help with the Musca infections, Amanda Manfredo for the RNA extractions and sequencing library preparations, and Dr. Richard Meisel and Dr. John Stoffolano for helpful comments and discussion. This work was supported by NIH grant R01 AI064950 to AGC and BPL.

## Supplemental Tables

Table S1. Species included in homology assignments, and data sources. (XLS)

Table S2. List of genes with annotated immune function in *D. melanogaster* (XLS)

Table S3. Full list of putative *M. domestica* immune-related genes (XLS)

Table S4. Gene ontology categories overrepresented in differentially regulated gene sets. (XLS)

Table S5. Gene duplication simulation input values and analyzed data.

